# Chimeric and Outward Reads in Short-Read RNA Sequencing Data: A Unified Circular RNA Assembly Framework

**DOI:** 10.64898/2026.11.15.765655

**Authors:** Anubhav Sen, Tasfia Zahin, Mingfu Shao

## Abstract

Circular RNAs (circRNAs) are RNA molecules that play important roles in regulating gene expression, yet detecting and assembling them from short-read RNA sequencing (RNA-seq) data remains challenging. Most methods rely on chimeric reads that cross the back-splicing junction (BSJ), but for lowly expressed circRNAs, these reads are often absent, so many circRNAs go undetected. In this work, we propose TERRACE 2.0, a method that addresses this limitation by leveraging an additional but underutilized signal: outward reads. These are paired-end reads that point away from each other when aligned to the reference genome, which is a direct consequence of sequencing a fragment from a circRNA. Together, this multi-signal framework enables circRNAs to be detected and assembled more accurately from short-read RNA-seq data. TERRACE 2.0 infers candidate BSJs from outward reads by testing which ones give a fragment length consistent with the library, reconstructs each full-length circRNA from a splice graph, and combines outward and chimeric reads in a single representation that selects among the candidates jointly. On eight human tissue samples, TERRACE 2.0 has the best average F-score with and without gene annotations. On an alignment that retains more chimeric reads, it is the most accurate assembler in every sample, while its predecessor TERRACE loses much of its precision. Most notably, outward reads are involved in 36% to 56% of the true circRNAs that TERRACE misses, and on simulated data TERRACE 2.0 also has the highest recall and F-score.

## 1 Introduction

Through alternative splicing, the exons of a single gene can be assembled in different arrangements to produce several distinct classes of RNA. The most common are linear transcripts (e.g., messenger RNA), in which exons are joined in order from the 5’ to the 3’ end of the gene. Some genes also produce a second class of RNA molecules in which exons are joined into a closed loop that lacks free ends. These molecules are known as circular RNAs (circRNAs). They arise when the spliceosome joins the 3’ end of a downstream exon with the 5’ end of an upstream exon, a process called back-splicing. CircRNAs were once regarded as rare by-products of splicing, but sequencing has shown that thousands of human genes produce them, often in patterns that differ between tissues. [10, 11, 15–18] Unlike linear transcripts, their “closed” form protects them from degradation by exonucleases. [10, 19] They have also been linked to conditions such as Alzheimer’s disease and certain cancers, making them promising candidates as biomarkers. [12, 14, 21, 23] Our work depends on first finding the circRNAs in a sample and determining the exons each is composed of. That is more challenging than it sounds; circRNAs are usually present at low levels and consist of the same exons as the much more abundant linear RNAs from the same gene. [7, 13, 20]

The hallmark feature that distinguishes a circRNA is its back-splicing junction (BSJ), which does not exist in linear transcripts. [9, 15, 20] BSJs are typically detected from chimeric reads, which are reads with supplementary alignments in which the 3’ end of the read maps upstream of its 5’ end. Tools such as find circ [15], CIRCexplorer2 [27], CIRI2 [6], and DCC [2] primarily report BSJs this way, and assemblers such as TERRACE [26] (the predecessor to this work, on which our method builds) and CIRI-full [29] use the detected BSJs as the starting point for reconstructing full-length circRNAs. [20, 22] This makes detecting the BSJ the critical first step. Due to the low expression levels of circRNAs, however, BSJs are often not sequenced at all, so a circRNA that is present in a sample can leave no chimeric read behind, and neither a detector nor an assembler has any way to find it if no such evidence exists. [7, 13, 22]

We recently discovered another strong indicator of circRNA expression in paired-end RNA sequencing (RNA-seq) data that has been largely overlooked: outward reads. In standard Illumina protocols, the two mates of a fragment normally point toward each other, which we refer to as inward reads. When a fragment comes from a circRNA, the mates can instead map in the reverse arrangement, so that they point away from each other on the genome. We refer to these as outward reads.

We propose TERRACE 2.0, a novel method for identifying BSJs and assembling circRNAs using outward reads. Essentially, the key idea is to infer the fragment whose length is consistent with the fragment length distribution. Specifically, let (*R*_1_, *R*_2_) be a pair of outward reads, and assume that the splice positions around the pair (the dashed vertical lines in Fig. 1, middle) have been identified. These positions can typically be inferred with high accuracy from the reads. [3] The circRNA from which (*R*_1_, *R*_2_) originates shares the same set of splice positions as its cognate linear RNA, so we examine whether the right splice position of a downstream exon of *R*_1_ can be connected to the left splice position of an upstream exon of *R*_2_ (Fig. 1, right). Once a BSJ is hypothesized, the length of the entire fragment can be calculated precisely. We estimate the median fragment length by sampling reads, discard hypotheses that imply a fragment much longer than the library produces, and keep how far each remaining fragment lies from the median as a measure of how well it fits.

**Fig. 1:**
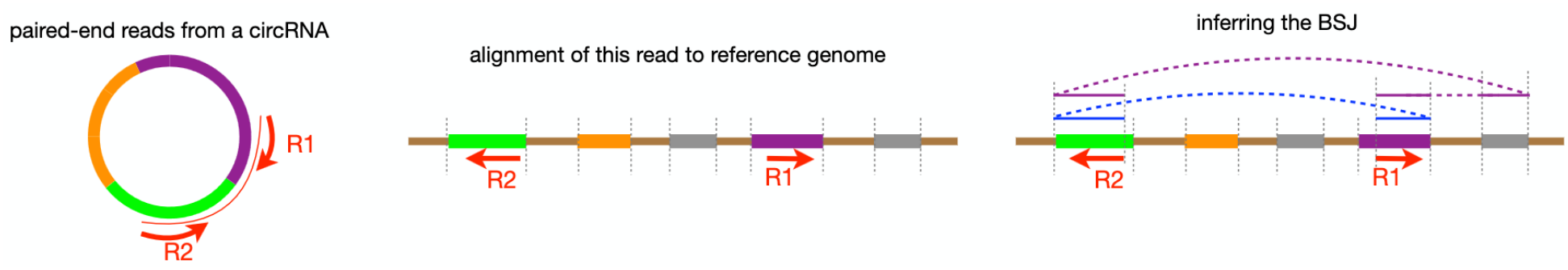
Detecting BSJs from outward reads. Left: two ends of a fragment from a circRNA get sequenced together. Middle: illustration of the alignment of *R*1 and *R*_2_ to the reference genome. They point in the opposite direction, a strong signal they originate from a circRNA. Note that neither end touches the BSJ. Right: inferring the BSJ from *R*_1_ and *R*_2_. The complete fragment can be either the blue one or the purple one, but the true one can be inferred based on the fragment length.

## 2 Methods

At a high level, TERRACE 2.0 takes a BAM file of paired-end total RNA-seq reads (along with the reference FASTA) and it produces a set of full-length circRNAs, each identified by its BSJ and its ordered list of exons. It is implemented as an extension of TERRACE [26] and works on one sample at a time. The reference sequence is used only for checking splice motifs and for recovering soft-clipped junctions.

### 2.1 Building the Splice Graph

We begin where TERRACE does. Reads on the same chromosome that overlap, directly or through a chain of overlaps, are grouped into a bundle, and each bundle is then processed independently. We retain the junctions supported by at least 10 reads and divide the bundle into non-overlapping regions at their splice sites and at the two ends of the bundle. These regions form the vertices of a splice graph *G* [8], and each one stores its start and end coordinates in the genome, along with its average coverage. Two vertices are joined by a directed edge when they are adjacent on the genome or when a junction connects them, and the weight of an edge signifies the number of reads that support it. Since every edge points from a smaller coordinate to a larger one, *G* is a directed acyclic graph whose vertices are already in topological order. Every read is then described by the ordered list of vertices it spans. In the same pass, we estimate the fragment length distribution by sampling reads and record its median, which we denote as *M*.

### 2.2 Evidence from Chimeric Reads

Evidence for a BSJ comes from the alignments, and the most direct kind comes from chimeric reads. Because a read that crosses a BSJ cannot be aligned in one piece, the aligner splits it into two alignments, and the “breakpoint” marks the BSJ itself. For each chimeric read, we gather its alignments by read name, order them by position, and take the BSJ coordinates from the outermost ends of the first and last alignments. When only a short segment of a read lies beyond the BSJ, the aligner usually soft-clips it instead. To recover these reads, we examine each read with a soft clip of at least 15 bases and look for a nearby splice position in the graph where the clipped sequence could belong. We then check whether the clip matches the reference sequence there, and we accept a match only when the Jaccard similarity is at least 0.9. Each accepted match is recorded as a new supplementary alignment and treated in the same way as those inferred from the alignment [26].

For every chimeric read, the sequence between the alignments is never observed directly, so we reconstruct it from the splice graph by finding paths that connect each alignment to the next. In Fig. 2, the read *R*_1_ is split by the BSJ into two alignments, *R*_1_.*A* and *R*_1_.*B*, and its mate *R*_2_ lies between them. Let *t*_1*A*_ be the last vertex of *R*_1_.*A*, let *s*_2_ and *t*_2_ be the first and last vertices of *R*_2_, and let *s*_1*B*_ be the first vertex of *R*_1_.*B*. Each gap then has known vertices at both ends, so *Path*_*A*_, from *t*_1*A*_ to *s*_2_, and *Path*_*B*_, from *t*_2_ to *s*_1*B*_, can be searched independently. A gap can be bridged by many different paths, and choosing one by a single criterion, such as the highest minimum edge weight, can be misleading, so we keep every path that is not dominated by another. Paths that reach the same vertex are compared on two criteria: the minimum edge weight along the path (higher is better) and the minimum vertex weight along the path (higher is better). A path is discarded if another is at least as good on both and strictly better on at least one. Because the splice graph is acyclic, this can be computed by dynamic programming in topological order. Joining the surviving paths across the gaps gives the full transcript, and each such combination is a candidate circRNA.

**Fig. 2:**
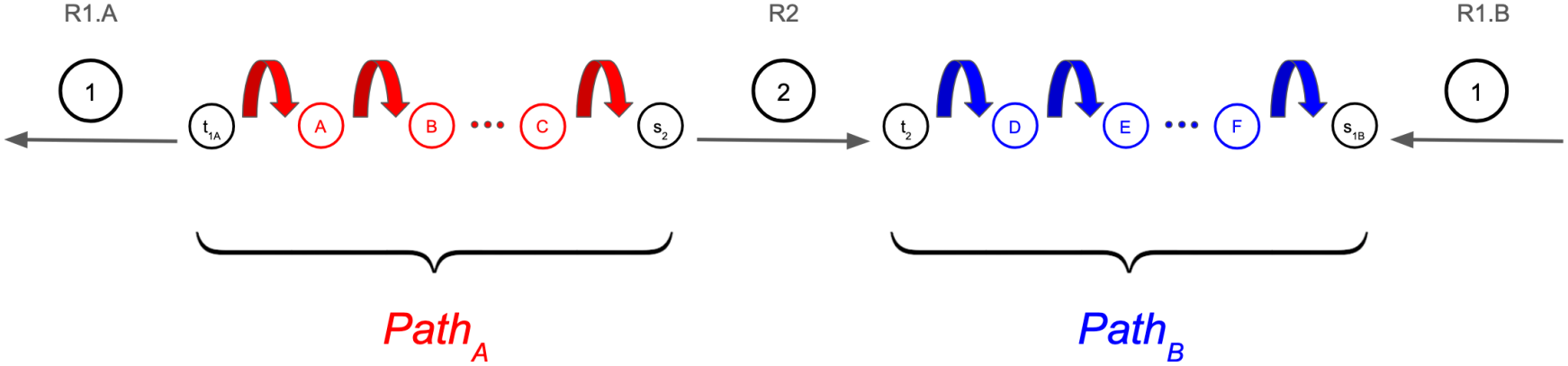
Reconstructing a circRNA from a chimeric read. Each gap between the alignments is bridged by a path through the splice graph, *Path*_*A*_ (red) and *Path*_*B*_ (blue), and each is one of many candidates. Together with the three alignments, the paths span the full circRNA. In this case, the BSJ is marked by where *R*_1_.*A* joins *R*_1_.*B*.

We can also reconstruct the fragment that a chimeric read came from. Using the example presented in Fig. 2, the fragment starts at the mate *R*_2_ and runs in the direction *R*_2_ maps. It spans all of *R*_1_ and crosses the BSJ, but it covers only one of the two gaps (Fig. 3): *Path*_*B*_ when *R*_2_ maps to the forward strand, and *Path*_*A*_ when it maps to the reverse strand. The fragment length is the sum of the lengths of the vertices it covers, so introns are not counted in the calculation. How far this length deviates from *M* is recorded and used later as a feature.

**Fig. 3:**
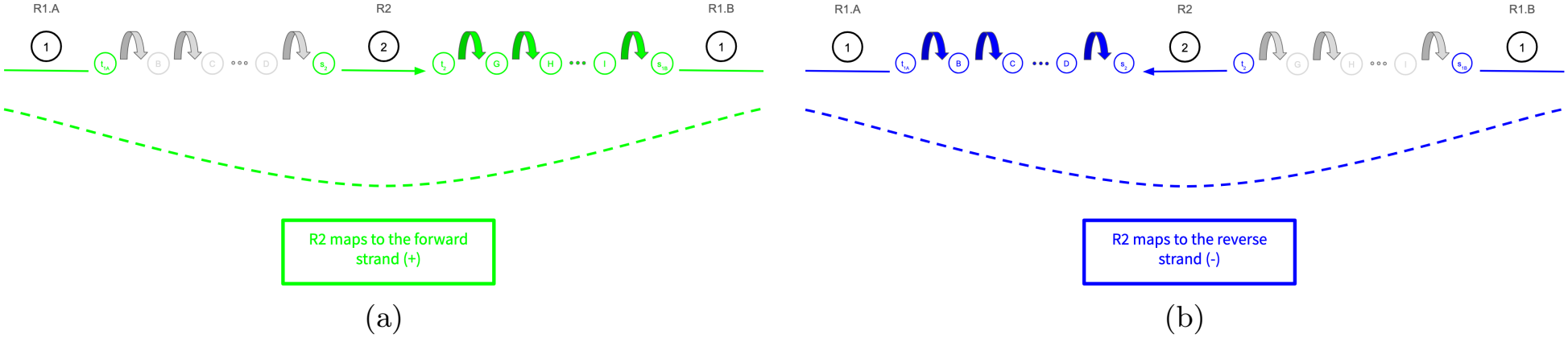
The fragment of a chimeric read. The fragment starts at *R*_2_, runs in the direction *R*_2_ maps, spans all of *R*_1_, and covers only one of the two gaps. (a) When *R*_2_ maps to the forward strand, the fragment covers *Path*_*B*_ but not *Path*_*A*_. (b) When *R*_2_ maps to the reverse strand, it covers *Path*_*A*_ but not *Path*_*B*_ . The fragment covers the parts of the circRNA highlighted in color. The BSJ is indicated by the dashed line.

### 2.3 Evidence from Outward Reads

The second type of evidence, outward reads, is what distinguishes our method from existing assemblers. Unlike a chimeric read, an outward read tells us that a BSJ exists but not where it is, so the next step is to hypothesize and pinpoint its location. As described in Fig. 4(a), let *R*_1_ be the upstream mate and *R*_2_ the downstream mate of the read pair. Each mate is described by the vertices of the splice graph that it covers. We let *s*_1_ and *t*_1_ denote the first and last vertices of *R*_1_, and *s*_2_ and *t*_2_ the first and last vertices of *R*_2_. If the pair comes from a circRNA, the fragment can be traced from *R*_2_ rightward to a splice site *R*, across the BSJ to a splice site *L* upstream of *R*_1_, and finally into *R*_1_. Let *v*_*L*_ be the vertex that starts at *L* and *v*_*R*_ the vertex that ends at *R*. Choosing *L* and *R* therefore fixes the BSJ, and what remains is to connect the known pieces into a full transcript with three paths through the graph: *Path*_*L*_ from *v*_*L*_ to *s*_1_, *Path*_*M*_ from *t*_1_ to *s*_2_, and *Path*_*R*_ from *t*_2_ to *v*_*R*_. The BSJ then joins *v*_*R*_ back to *v*_*L*_. Candidates for *L* are the splice sites at or before the start of *R*_1_, and candidates for *R* are the splice sites at or after the end of *R*_2_. For each candidate *L* we enumerate the possible *Path*_*L*_, for each candidate *R* we enumerate the possible *Path*_*R*_, and we enumerate *Path*_*M*_ between *t*_1_ and *s*_2_. As in the chimeric case, each of the three searches keeps only non-dominated paths. We then combine the surviving paths from the three searches.

**Fig. 4:**
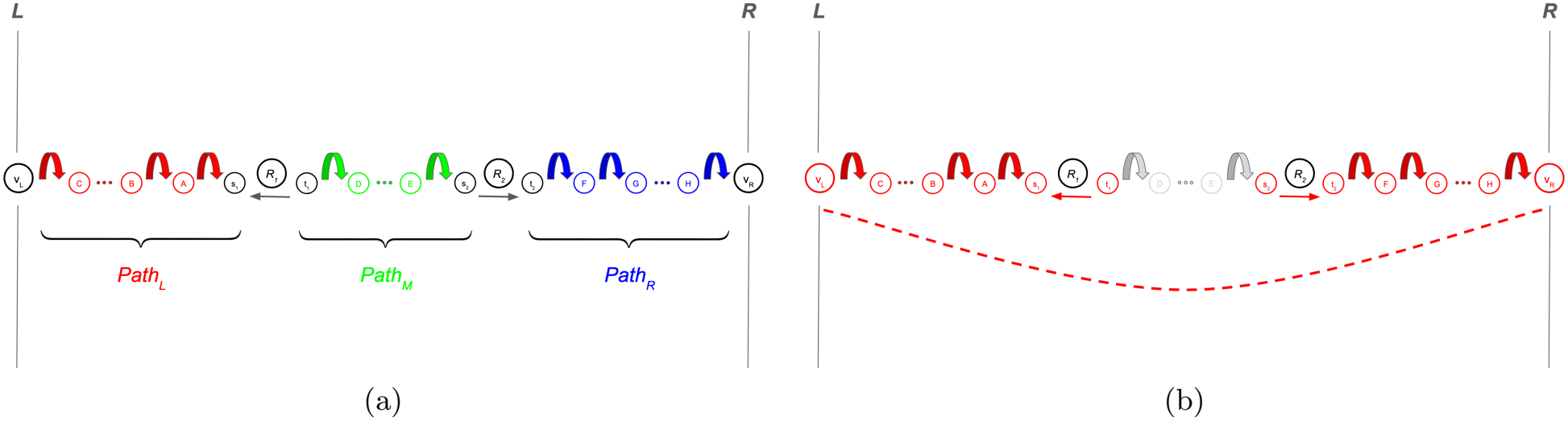
(a) Reconstructing a circRNA from an outward read. The mates *R*_1_ and *R*_2_ point away from each other. *Path*_*L*_ (red) connects *L* to *R*_1_ through the splice graph, *Path*_*M*_ (green) connects *R*_1_ to *R*_2_, and *Path*_*R*_ (blue) connects *R*_2_ to *R*. Together with the two mates, the three paths span the full circRNA transcript, and the BSJ is the point where *R* joins *L*. (b) The fragment implied by a hypothesized BSJ for an outward read. The fragment covers the parts of the circRNA highlighted in red. Note that *Path*_*M*_ (grey) does not contribute to the fragment length. The BSJ is shown as the dashed line.

Each combination of *Path*_*L*_ and *Path*_*R*_ is now a hypothesis about the BSJ, and the library lets us test it. Because the positions of the mates and the splice sites are known, we can compute exactly how long the fragment would have been if the hypothesis were true. As shown in Fig. 4(b), a fragment inferred from an outward read covers the circRNA from *v*_*L*_ through *R*_1_ on one side of the BSJ, and from *R*_2_ through *v*_*R*_ on the other. The fragment length is the sum of the lengths of the vertices on these two stretches, so introns are not counted. We keep a combination only if its fragment length is at most 2*M* and record its deviation from *M*, which the subsequent scoring steps use as a feature. Note that the fragment length does not depend on *Path*_*M*_, so the test is run once for each pair of *Path*_*L*_ and *Path*_*R*_ and applies to every *Path*_*M*_ . Each accepted combination, joined with *Path*_*M*_, is a candidate circRNA supported by the outward read.

### 2.4 Combining the Evidence

We now have two sets of candidates, one built from chimeric reads and one from outward reads. We combine them in a single bipartite graph representation with reads on one side and candidates on the other. A read is connected by an edge to every candidate that was built from its alignments. A candidate can therefore be supported by chimeric reads, outward reads, or both. A read may also support several competing candidates, which is expected at loci containing more than one isoform. A read supporting several candidates connects them all, so the graph decomposes into groups of candidates connected through shared reads. We call each group a connected component (CC) of this graph.

Each candidate is then described by a set of features, including the number of chimeric and outward reads that support it, the minimum edge and vertex weights along its path, the number and lengths of its exons, and how far its fragment length deviates from *M* . We then reduce the pool of candidates in two ways. First, the same BSJ may appear as several nearly identical candidates. This happens because of microhomology. When a short identical sequence occurs at both ends of a junction, the junction can be placed at more than one position. To handle this edge case, we place candidates whose BSJs lie within one read length of each other (150 bases in our data) in the same CC, even when they share no reads. Second, we drop a candidate *X* when another candidate *Y* is better supported on every count: all the reads that support *X* also support *Y*, *Y* has a higher minimum edge weight, and its fragment length lies closer to *M*.

### 2.5 Selecting circRNAs

The pool that remains is still large and mostly false, so we first filter it with a random forest of 30 trees with depth 10 and balanced class weights, which we call the Pre-Filter. It is trained on the candidate features, and because dropping a true candidate at this stage cannot be undone, we choose the probability cutoff to remove as many false candidates as possible while keeping at least 98% of the true ones in cross-validation.

What remains is chosen by a greedy procedure that runs separately on each CC. Within a CC, every read starts out as “unexplained” and becomes marked as “explained” once a circRNA claims it, so a read counts toward only one circRNA and the procedure avoids reporting many near-identical isoforms. In each round, a random forest trained on the candidate features, which we call the Selector, scores every remaining candidate using only the unexplained reads, and the best-scoring candidate is selected. If it passes a set of validity checks (described in the following paragraph), it is output and its reads are marked as explained. If it fails, it is discarded, and its reads remain available to the other candidates. The procedure ends when all reads have been claimed by a circRNA.

A candidate selected in a round is output only if it passes four stringent checks. The first is a minimum evidence check. The candidate must have at least two supporting reads, as candidates supported by a single read were almost always false in our data. The second is a splice motif check. Every splice junction in the transcript, including the BSJ, must show the canonical GT-AG splice motif (or CT-AC on the reverse strand), the motif commonly found at the vast majority of splice junctions. [1] The third is a structural check. A candidate may have at most 15 exons, and its exons may be at most 2,000 bases long for a single-exon circRNA and at most 1,000 bases long for a multi-exon circRNA. The fourth restricts the output to the primary chromosomes (chr1 to chr22, chrX, and chrY in humans).

### 2.6 Recovering Missed circRNAs

A true circRNA can still be lost during greedy selection, for instance when a sibling isoform wins first and consumes the reads that the two share. To recover some of these cases, we make a last pass over every candidate that was never output, once all CCs in the bundle are finished processing. This pass uses a third model, which we call the Recoverer, a calibrated gradient-boosted tree model that estimates the probability that each candidate is genuine from the full feature set. The full list of features, and which of the three models (the Pre-Filter, Selector, and Recoverer) use each of them, is provided in Appendix A. Beyond the candidate features described above, the Recoverer also uses the history of each candidate during selection (for example, how many rounds it competed in and which check rejected it), its relation to the circRNAs already output for the bundle (for example, whether it repeats one of them or shares one end with it), and the splice motif at each end of its BSJ. A candidate is added when the probability is at least 0.5, so there is no fitted cutoff. This pass only adds to the output and never changes an earlier decision.

## 3 Results

We evaluated TERRACE 2.0 both on simulated and real data. The real dataset is taken from eight human tissue samples: brain, heart, kidney, liver, lung, prostate, skeletal muscle, and testis collected from a long-reads circRNA study [30]. This work also catalogs full-length circRNAs assembled from long reads and gene annotation that we use as ground truth. We compare TERRACE 2.0 with four existing assemblers: CircAST [24], CIRCexplorer2 [27], CIRI-full [29], and TERRACE [26]. These tools differ in whether they need a gene annotation. CIRCexplorer2 and CircAST cannot run without one, whereas CIRI-full, TERRACE, and TERRACE 2.0 can run with or without it. The distinction matters because an annotation may be incomplete for a given sample, or unavailable altogether. We therefore report two sets of results. In the first, no annotation is provided, so TERRACE 2.0 is compared with CIRI-full and TERRACE. In the second, every tool receives the same annotation, which allows a comparison with all four existing assemblers.

We compared all five assemblers by precision, recall, and F-score against a ground truth of full-length circRNAs in the same samples from [30], counting a prediction as correct only when both its BSJ and its exon chain match. This ground truth may miss some circRNAs that are truly expressed, so precision may be underestimated while still allowing comparisons of relative performance across tools. Since a single output can hide the trade-off between precision and recall, we also drew a precision-recall curve for each framework by sweeping the threshold on its confidence score. When no framework is best on both measures, we compared their precision at the same recall.

TERRACE and TERRACE 2.0 start from an aligned BAM file, so their results can depend on how the reads were aligned. We therefore ran both of them on two different alignments of the same reads, which we describe next.

### 3.1 Regular and Relaxed Alignments

We aligned the same reads with STAR [3] in two separate ways, which we call the “regular” alignment and the “relaxed” alignment. They differ in how strictly STAR filters a read that aligns partially. In the relaxed alignment, we lower the minimum fraction of a read that must align from STAR’s default of 66% to 50%, by setting both `--outFilterScoreMinOverLread` and `--outFilterMatchNminOverLread` to 0.5. Chimeric reads benefit most from this change. A read that crosses a BSJ aligns to two different places, so only a fraction of it aligns at each place, and a lower threshold is more likely to keep it. The relaxed alignment therefore preserves more chimeric reads and gives an assembler more BSJs to work with, but it also contains more spurious alignments. CIRI-full does not use our alignments while CIRCexplorer2 and CircAST are unaffected by the relaxed setting, so their results are the same under both.

### 3.2 Regular Alignment

Without annotations, TERRACE 2.0 achieves the highest F-score in seven of the eight samples (Fig. 5 and Table 1). The gain over TERRACE is modest, 1.18 points on average. It is largest in heart, brain, and testis, and TERRACE is slightly ahead in prostate (26.87% against 26.52%). The two assemblers reach a similar average precision, 42.65% for TERRACE and 41.58% for TERRACE 2.0, so the difference comes from recall, which averages 18.88% for TERRACE 2.0 and 17.72% for TERRACE. CIRI-full is far behind both.

**Table 1:** Average F-score (%) across the eight samples for each framework, on the regular and relaxed alignments, without and with annotations. A dash means that the tool cannot run without an annotation. The results of CircAST, CIRCexplorer2, and CIRI-full do not depend on the alignment.

| Framework | Regular alignment |  | Relaxed alignment |  |
|---|---|---|---|---|
|  | w/o anno. | with anno. | w/o anno. | with anno. |
| CircAST | - | 5.28 | - | 5.28 |
| CIRCexplorer2 | - | 20.77 | - | 20.77 |
| CIRI-full | 10.45 | 10.35 | 10.45 | 10.35 |
| TERRACE | 22.49 | 23.43 | 18.24 | 18.72 |
| TERRACE 2.0 | 23.67 | 23.80 | 24.50 | 24.70 |

**Fig. 5:**
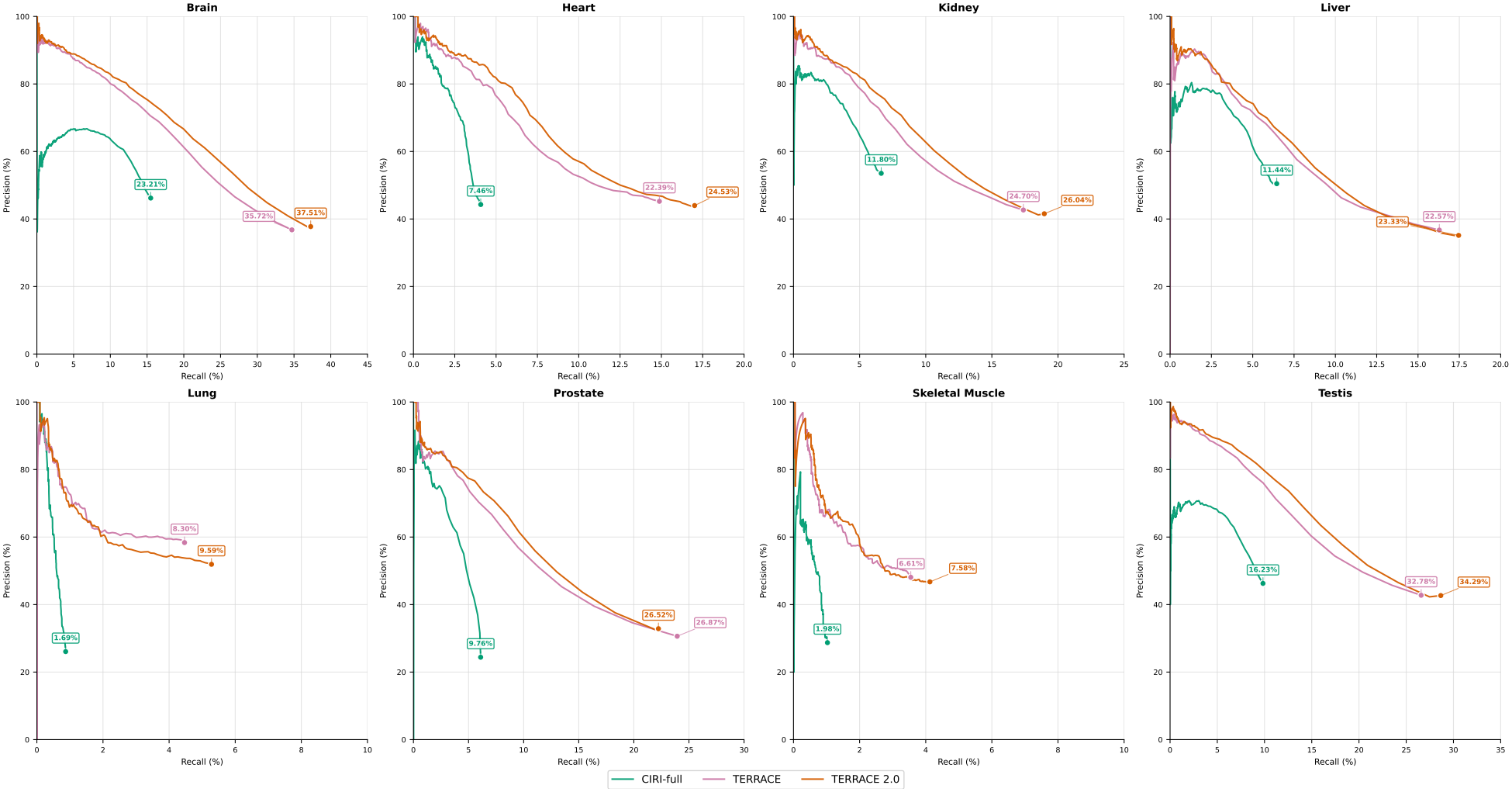
How the frameworks compare in assembly accuracy on the regular alignment when annotations are not provided. F-scores are shown next to the end of each curve.

With annotations, TERRACE 2.0 still has the highest average F-score (Fig. 6 and Table 1). CIRCexplorer2 is the strongest of the tools that depend on the annotation, and both TERRACE and TERRACE 2.0 outper-form it in every tissue, by 2.66 and 3.03 points, respectively, on average. The annotation helps TERRACE more than TERRACE 2.0, so the gap between the two narrows to 0.37 points. TERRACE is now ahead in lung, prostate, and skeletal muscle, by 0.34 to 1.12 points, and TERRACE 2.0 leads in the other five tissues.

**Fig. 6:**
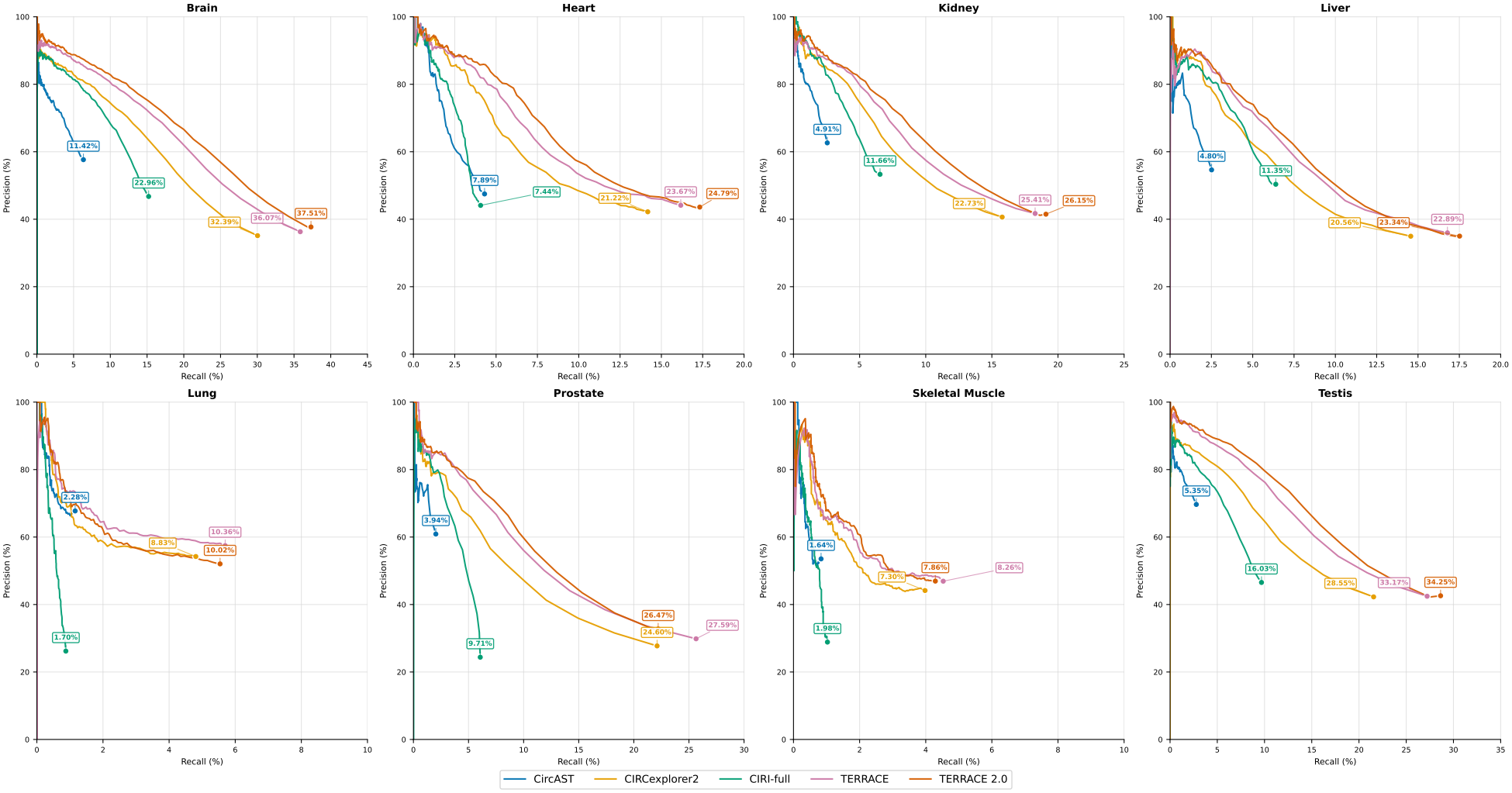
How the frameworks compare in assembly accuracy on the regular alignment when annotations are provided. F-scores are shown next to the end of each curve.

### 3.3 Relaxed Alignment

The relaxed alignment separates the two assemblers much more clearly. Without annotations, TERRACE 2.0 achieves the highest F-score in all eight samples (Fig. 7). Its average F-score is 24.50%, compared with 18.24% for TERRACE, and the gain over TERRACE is positive in every sample. It is largest in the prostate where the F-score rises from 10.51% to 27.33%. It is smallest in the two tissues with the fewest detectable circRNAs, lung (8.68% to 10.63%) and skeletal muscle (7.25% to 8.52%), where every tool recovers only a small fraction of the ground truth circRNAs.

**Fig. 7:**
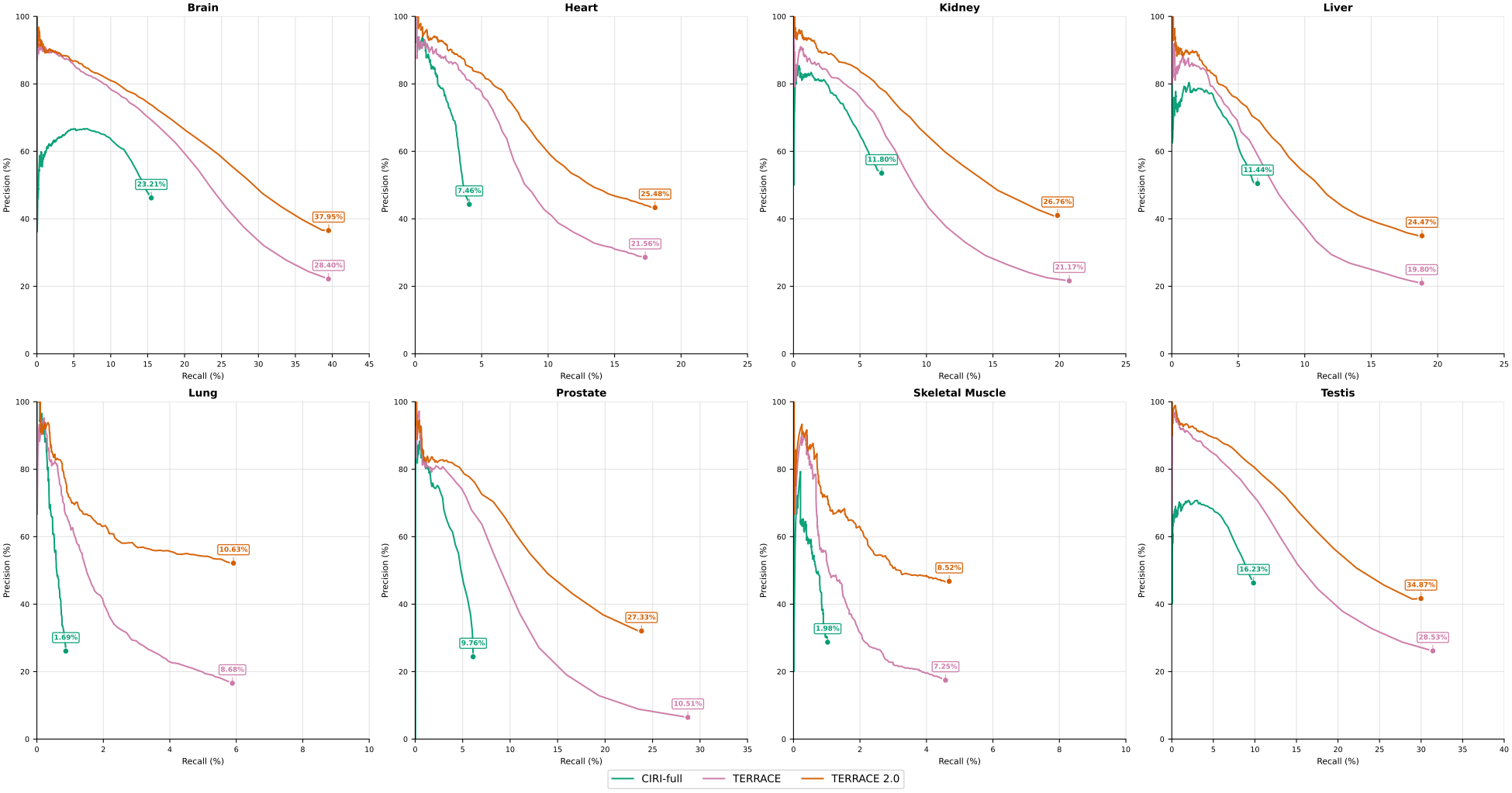
How the frameworks compare in assembly accuracy on the relaxed alignment when annotations are not provided. F-scores are shown next to the end of each curve.

The endpoints of the curves in Fig. 7 show where the gain comes from. TERRACE 2.0 reaches an average precision of 41.05%, more than double the 19.98% of TERRACE, and it is more precise in every tissue. The effect is strongest in prostate, where TERRACE reports many predictions that match no ground truth circRNA and reaches a precision of only 6.43%. This costs little recall, which averages 20.08% for TERRACE 2.0 and 20.87% for TERRACE. Recall is lower in only three samples, kidney, testis, and prostate (by 0.88 to 4.88 points), and in every tissue the precision gained is much larger than the recall given up. In other words, TERRACE 2.0 reaches essentially the same recall as TERRACE with roughly half as many predictions, and nearly all of the predictions it drops are false.

With annotations, TERRACE 2.0 again has the highest F-score in all eight samples (Fig. 8). It outperforms CIRCexplorer2, the strongest tool that depends on the annotation, by 3.93 points on average (24.70% against 20.77%), with a margin that ranges from 1.75 points in skeletal muscle to 6.29 points in testis. The advantage comes mostly from recall (20.26% against 15.88%), at a similar precision (41.00% against 40.16%).

**Fig. 8:**
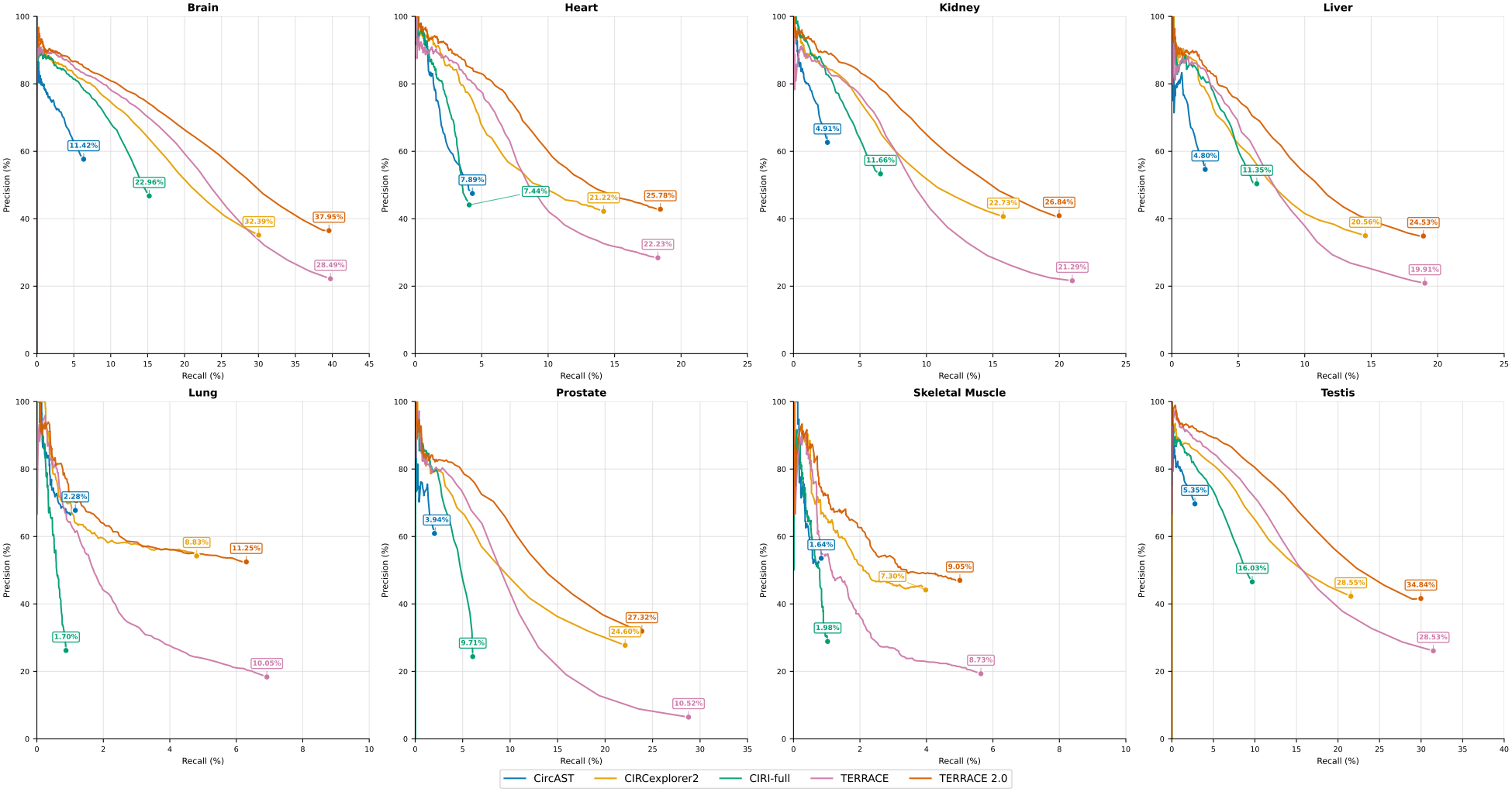
How the frameworks compare in assembly accuracy on the relaxed alignment when annotations are provided. F-scores are shown next to the end of each curve.

Table 1 puts the two alignments side by side and explains why the relaxed alignment favors TERRACE 2.0. Relaxing the alignment gives both assemblers more recall. For TERRACE, the average recall rises from 17.72% to 20.87%, but its average precision falls from 42.65% to 19.98%, so its average F-score drops from 22.49% to 18.24%, with a fall in six of the eight samples. The extra chimeric reads include many spurious ones, and TERRACE has no way to distinguish them. In contrast, TERRACE 2.0 takes in the same reads without losing precision (41.58% to 41.05%), and its F-score improves in every sample, from 23.67% to 24.50% on average. It is therefore the assembler that can make use of the additional reads, and its best result comes from the relaxed alignment.

### 3.4 What Outward Reads Contribute

Since the outward reads are the key feature that distinguishes TERRACE 2.0 from TERRACE, we asked how many of the circRNAs it detects depend on them. We divided the detected circRNAs according to the reads that support each one: chimeric reads only, outward reads only, or both (Fig. 9). In all four settings, most circRNAs belong to the first group. About 76% of the circRNAs detected on the regular alignment and 77% of those detected on the relaxed alignment are supported by chimeric reads alone, 15% to 16% by both kinds, and 7% to 8% by outward reads alone.

**Fig. 9:**
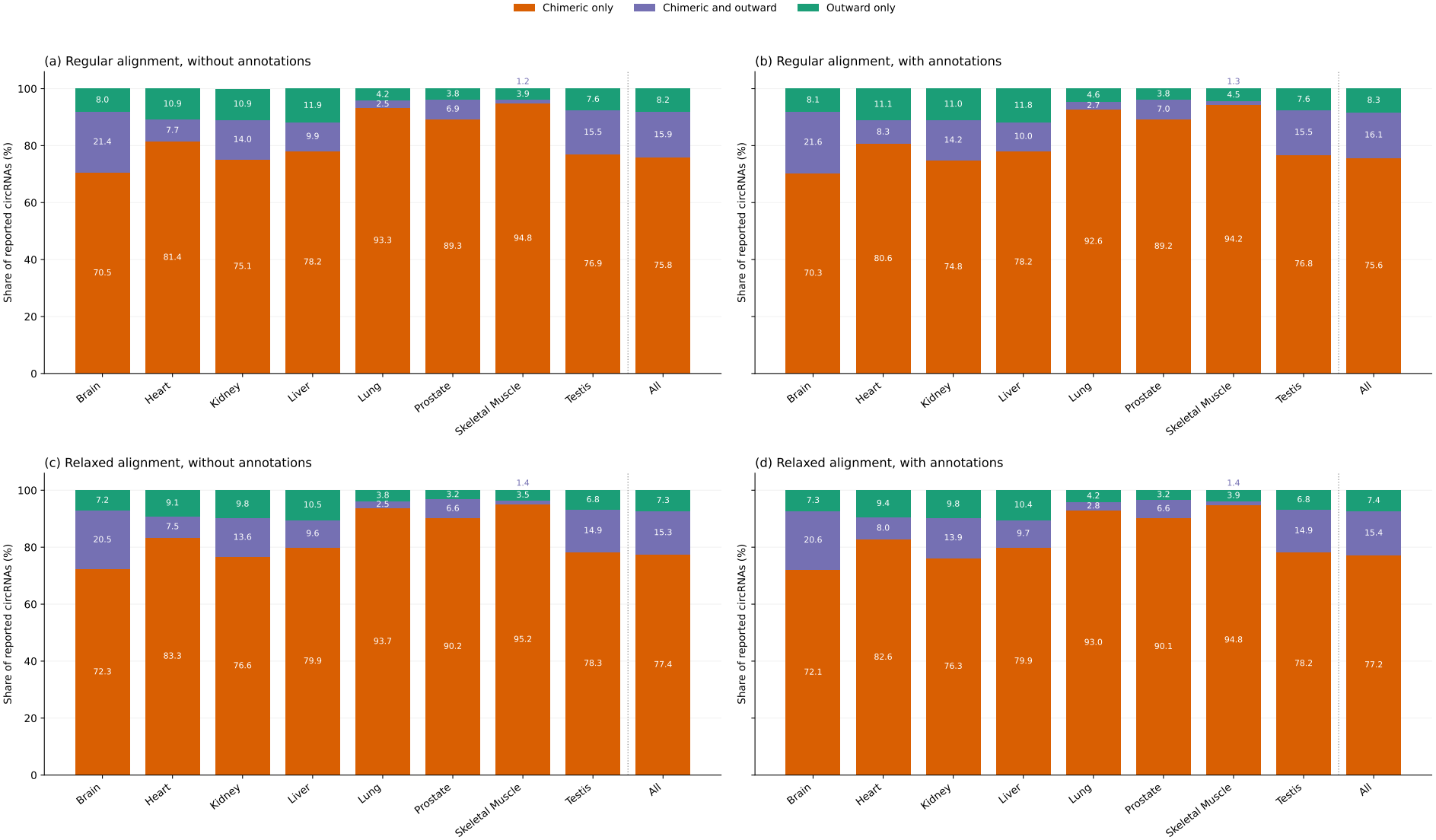
Breakdown of the circRNAs reported by TERRACE 2.0 by supporting reads (chimeric reads only, outward reads only, or both) in each sample and pooled over all eight, on (a) the regular alignment without annotations, (b) the regular alignment with annotations, (c) the relaxed alignment without annotations, and (d) the relaxed alignment with annotations. The number on each segment is its percentage of the reported circRNAs, and the label of the thin segment for skeletal muscle (both kinds of support) is placed above its bar.

The groups differ sharply in how often they are accurate. Without annotations, about 65% to 67% of the circRNAs supported by both kinds of reads match the ground truth, compared with about 36% to 37% of those supported by chimeric reads alone, so when an outward read agrees with a BSJ that chimeric reads already point to, the prediction is far more likely to be correct. This group yields about a quarter of all true positives, but its size depends on the tissue (Fig. 9). It makes up about 21% of the circRNAs detected in brain and 15% in testis, and under 3% in lung and about 1% in skeletal muscle, the two samples in which every tool recovers the fewest circRNAs.

The outward-only group is where the outward reads stand alone. These circRNAs have no chimeric read across their BSJ, and nearly all of them (1,316 on the regular alignment and 1,197 on the relaxed one, without annotations) are indeed missed by TERRACE. There are 1,332 and 1,202 true positives of this kind, about 3% to 4% of all true positives. The group is small, between 3% and 12% of the circRNAs detected in a sample, and it comes at a price. Because no read crosses the BSJ, these are the hardest candidates to verify, and only about 16% to 17% of its predictions are correct, which is why TERRACE 2.0 asks for more support before it reports them (Section 2.5).

Together, the two groups show how much the outward reads contribute to what TERRACE 2.0 gains over TERRACE. Without annotations, TERRACE 2.0 detects 5,670 true circRNAs that TERRACE misses on the regular alignment, which we call “novel,” and 2,039 of them (36%) have outward support, either alone or together with chimeric reads. On the relaxed alignment, it detects 3,144 novel circRNAs, and 1,768 (56%) have outward support.

### 3.5 Simulated Data Runs

On real data, recall and precision are measured against a ground truth that may not list every circRNA that is expressed. Simulated data removes this uncertainty, so we also ran all of the tools on the existing simulated datasets [5, 26]. We use the configuration with read length 100, circular read coverage 10, and linear read coverage 10, and we report the mean and standard deviation over ten replicates (Fig. 10). As with the real data, TERRACE and TERRACE 2.0 were run on both the regular and the relaxed alignment, and every tool was run without and with annotations.

**Fig. 10:**
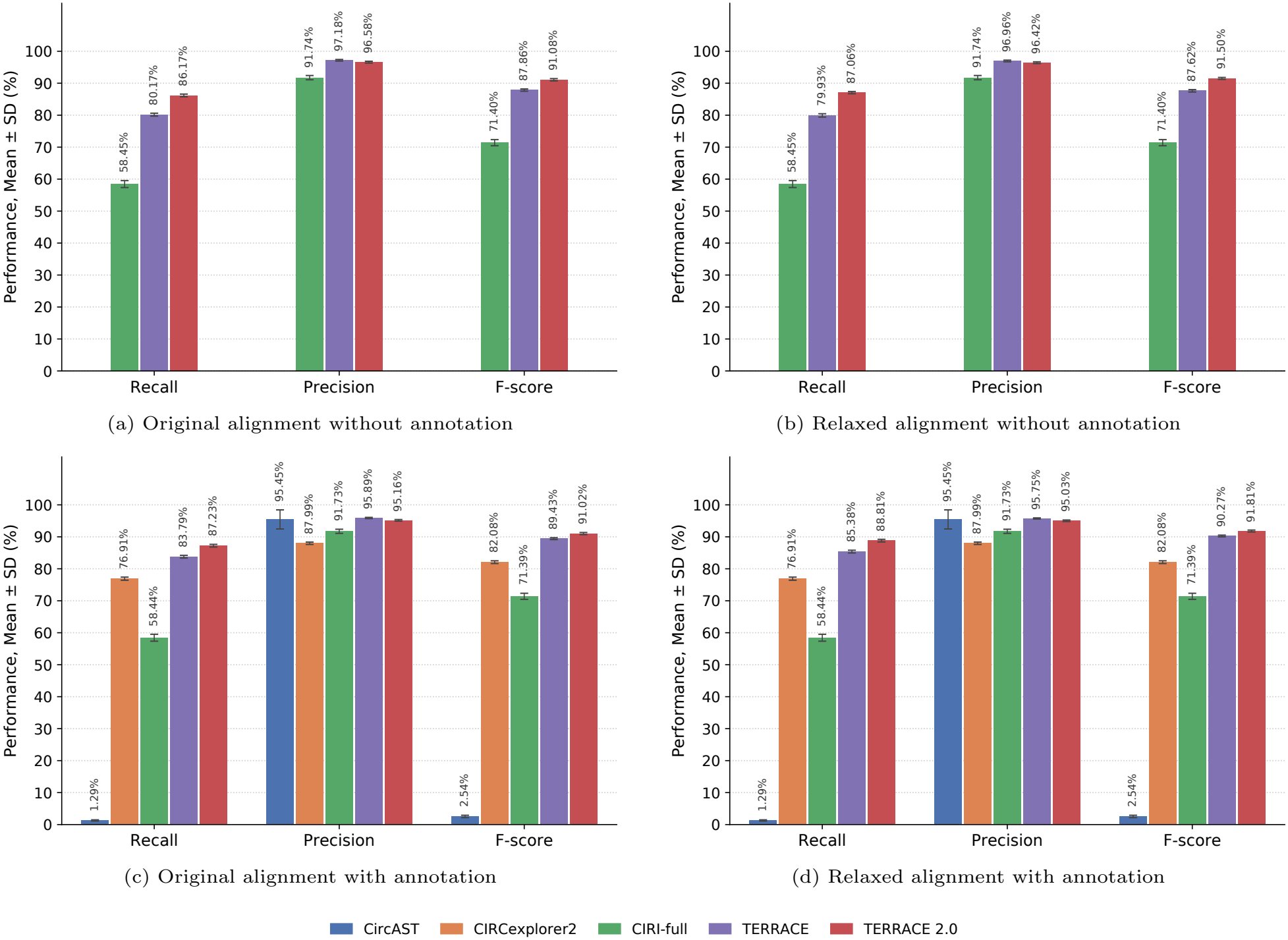
Comparison of simulation results for two alignment settings, with and without annotation. Error bars show variability across 10 samples.

TERRACE 2.0 has the highest recall and F-score in all four settings. Without annotations on the regular alignment, its F-score is 91.08%, compared with 87.86% for TERRACE and 71.40% for CIRI-full. The gain comes from recall, which is 86.17% for TERRACE 2.0 and 80.17% for TERRACE, at a nearly identical precision (96.58% against 97.18%). The relaxed alignment gives the same picture (F-score 91.50% against 87.62%, recall 87.06% against 79.93%), and so do the runs with annotations, where the F-score of TERRACE 2.0 is 91.02% and 91.81% on the two alignments, against 89.43% and 90.27% for TERRACE and 82.08% for CIRCexplorer2. CircAST, as on the real data, recovers almost nothing (recall 1.29%). The precision of TERRACE 2.0 also remains above 95% in every setting, so its higher recall does not come at the cost of more false calls.

## 4 Conclusion and Next Steps

Short-read circRNA assemblers have so far depended on a single kind of evidence, chimeric reads, which are uncommon for lowly expressed circRNAs. TERRACE 2.0 shows that this is not the only evidence available. By testing which BSJs would give a fragment length consistent with the library, it can deduce a BSJ from outward reads even when no read crosses it. By combining outward and chimeric reads on a single graph in which candidates compete for the reads they claim, it keeps a circRNA only if it explains reads that no better candidate already explains. As a result, TERRACE 2.0 finds circRNAs that earlier methods miss without flooding its output with false predictions. It has the best average F-score in all four settings we tested, it is the most accurate assembler in every sample on the relaxed alignment, and it is far less sensitive than TERRACE to how the reads were aligned. Its accuracy also barely depends on an annotation, so it can be used for species and samples whose transcriptomes are poorly annotated. Most importantly, outward reads are also involved in between 36% and 56% of the true circRNAs that TERRACE misses.

Recall is still modest for every method on real data, and a BSJ that no fragment in the library points to cannot be recovered by any method. Some limits are also built into our design. The structural checks and the rule that keeps an outward read only if its fragment length is at most 2*M* are fixed, so a circRNA that exceeds them is dropped however well it is supported. The most direct improvement would be to replace these rules with a likelihood under the estimated fragment length distribution, or with limits learned from the data. Alignment offers a second route. Since the relaxed alignment helps TERRACE 2.0, a strategy that keeps chimeric reads while limiting spurious ones may raise recall further. A single model trained on all of the signals could also replace the separate stages and fixed checks. Beyond these refinements, there is room to broaden the framework. It should, of course, be tested on other samples and species to confirm that the gains hold beyond what we tested. A natural next step is to go beyond finding circRNAs and measure them, by estimating the abundance of each isoform and comparing it across tissues. Overall, TERRACE 2.0 is a first step toward capturing the information that outward reads carry, and we expect the circRNAs it reveals in existing data to be the beginning of a more complete picture.

## Availability

The source code of TERRACE 2.0 is freely available at https://github.com/Shao-Group/TERRACE/outward. The raw sequencing (FASTQ) files of the eight samples is available at BIGD (accession number: PR-JCA000751).

## Acknowledgment

This work is supported by the US National Science Foundation (2019797 and 2145171 to M.S.) and by the US National Institutes of Health (R01HG011065 to M.S.). We thank Carl Zang for his helpful input.

## Appendix A

Three models score candidates in TERRACE 2.0: the Pre-Filter, Selector, and Recoverer. They do not all use identical features, but they draw on the same pool. The Pre-Filter sees every feature that describes a candidate on its own (groups 1 to 6 below). The Selector uses a smaller set from the same groups; instead of the total number of supporting reads, it counts only the reads that are still unexplained, so a candidate’s score drops as better, first-picked candidates claim its reads. The Recoverer uses all of these features, along with two more groups (7 and 8) that can only be computed once selection has finished.

1. Read support. These are the number of chimeric reads and number of outward reads that support the candidate. We also count the distinct reads after merging reads that start and end at the same position, which are likely PCR duplicates. We observed that false candidates tend to be supported by more duplicated reads than true candidates. Duplicates also occur naturally in RNA-seq, so a high count alone is not enough to reject a candidate. We therefore give the count to the models as a feature and do not use it as a hard rule.
2. Quality of the chimeric reads. For each candidate, we record how many of its chimeric reads were recovered from a soft clip (a “fake” supplementary alignment) instead of being split by the aligner, the longest matched stretch of a supplementary alignment, and the anchor length (which is the shorter of the two aligned stretches on either side of the BSJ). A longer anchor means more of the read aligns reliably on each side of the BSJ, so the BSJ is more likely to be placed correctly.
3. Support along the path. We record the minimum edge weight and the minimum vertex weight along the path. Each of these reflects only the weakest point, so we add two counts that capture how much of the path has weak or no support: the number of edges with weight below the 10-read cutoff for keeping a junction, and the number of vertices that no read covers. The second count matters because the minimum vertex weight ignores uncovered vertices, and a path that crosses an unsupported region could otherwise appear well supported.
4. Fragment length. This is the fragment length implied by the candidate and its deviation from *M* . The deviation is the feature that tests how well the candidate agrees with the library. It is computed from the fragments reconstructed from the candidate’s supporting reads, whether they are chimeric reads, outward reads, or both (Sections 2.2 and 2.3).
5. Structure and local coverage. These include the number of vertices and exons, the total, maximum, minimum, and average exon length, its length on the genome (including introns), and the fraction of that length that is exonic. These structure features help because false candidates often have an unusual shape, such as a large number of exons or exons that are unusually long, especially in highly expressed regions. We also include the ratio of the candidate’s chimeric support to the highest coverage in its bundle. Highly expressed genes produce many noisy reads, so the same number of chimeric reads is less convincing there than in a “quiet” region, and the ratio puts the candidate’s support in that context.
6. Splice motifs. These features describe the sequence around the splice sites of a candidate. For the BSJ, we record the two bases at each end and whether they form a canonical motif, which is AG followed by GT on the forward strand and AC followed by CT on the reverse strand. For the other splice junctions in the transcript, we record the fraction that are not canonical. We also count inverted repeats, which are short stretches on the two sides of the BSJ that are reverse complements of each other. Because these repeats can pair with each other and bring the splice sites together, they serve as a signal that favors back-splicing, and candidates with more of them are more likely to be genuine circRNAs. [10, 28] The following two groups are used only by the Recoverer, since they primarily describe what happened to a candidate during selection.
7. Competition history. These are the number of rounds in which the candidate competed, the earliest and latest round it appeared in, whether it ever won a round, its best margin against the winner, the largest number of unexplained chimeric and outward reads it had in any round, and which of the four validity checks (described in Section 2.5) rejected it. The Recoverer needs to know how close a rejected candidate came to being chosen. A narrow loss, or a failure on a single check, suggests a real circRNA that was lost during selection, whereas a large gap to the winner in every round does not.
8. Overlap with output circRNAs. These features compare a candidate with the circRNAs already output for its bundle. Some show whether the candidate repeats one of them, by recording whether an output circRNA has the same BSJ, the same start, or the same end. Others describe the output around the candidate: how many circRNAs the bundle has, whether any of them overlaps the candidate on the genome, and how close the nearest one is. Finally, one feature compares the candidate with its rivals, recording where it ranks by supporting reads among all candidates with the same BSJ. A candidate that only repeats an isoform already output is usually a copy, but one that shares a BSJ and differs in its exons may be a real alternative isoform.

## Appendix B

**Table 2:** Number of chimeric reads (second column) and outward reads (third column) in each of the eight total RNA-seq samples. Note how outward reads are common in every sample, ranging from about a third as many as chimeric reads in lung to nearly as many in heart and liver.

| Sample | Number of Chimeric Reads | Number of Outward Reads |
|---|---|---|
| Brain | 707,852 | 382,805 |
| Heart | 358,780 | 353,395 |
| Liver | 230,789 | 216,047 |
| Lung | 1,387,404 | 447,880 |
| Kidney | 340,610 | 274,628 |
| Muscle | 734,215 | 266,586 |
| Prostate | 948,853 | 379,854 |
| Testis | 441,682 | 290,711 |

